# Cryo-ET and Sub-Volume Analysis of Fibrillar Pathology in the Neurodegenerative Human Brain

**DOI:** 10.64898/2026.09.04.749518

**Authors:** Lukas van den Heuvel, Daniel A. Stähli, Julika Radecke, Jean Daraspe, Notash Shafiei, Amanda J. Lewis, Wilma D. J. van de Berg, Christel Genoud, Anne-Laure Mahul-Mellier, Henning Stahlberg, Wen-Lu Chung

## Abstract

Structural studies of amyloid fibrils extracted from brain tissue have identified disease-specific fibril polymorphs. However, the mechanisms driving distinct polymorphs remain unclear because no method currently links the cellular context, composition and ultrastructure of individual aggregates to their constituent fibril polymorphs. Here, we present a workflow for *in situ* cryo-electron tomography (cryo-ET) to study amyloids within individual aggregates in the human neurodegenerative postmortem brain. Using chemically fixed tissue, we studied *in situ* tau fibrils in the hippocampus of an Alzheimer’s disease (AD) donor and α-synuclein fibrils within a glial nuclear inclusion in the cingulate gyrus of a multiple systems atrophy (MSA) donor. Subtomogram averaging and helical reconstruction of fibril subvolumes yielded low-resolution (∼30-35Å) density maps, which were compared to existing *ex vivo* structures. Preserved ultrastructure enabled detailed observations of pathology, including an α-synuclein fibril penetrating the nuclear envelope, providing mechanistic insight into intranuclear aggregate formation. Combined with existing high-resolution fibril structures lacking spatial context, cryo-ET offers a powerful approach to understanding amyloid polymorphism in neurodegenerative diseases, advancing anti-amyloid therapies and diagnostic tools.

## Main

Neurodegenerative diseases are a growing global health crisis, affecting over a hundred million people worldwide^1^. Most neurodegenerative diseases are associated with the misfolding of proteins and their accumulation into amyloid fibrils in the brain. Given the prion-like transmission properties of amyloid fibrils and their toxicity, they are believed to play a causative role in the spread of pathology and progressive neurodegeneration^2,3^. Understanding how physiological proteins adopt pathogenic conformations is therefore critical to understanding disease progression and developing disease-modifying therapies.

The study of the atomic structure of amyloid fibrils from the diseased brain became possible after the development of biochemical extraction methods isolating the fibrils from brain tissue, as well as cryo-electron microscopy (cryo-EM) structure determination and helical reconstruction^4^. The most effective extraction method isolates amyloid fibrils from bulk homogenates of ∼500mg tissue, based on their insolubility in the detergent sarkosyl^5^. Combining sarkosyl extraction with cryo-EM and single particle analysis (SPA) has led to the identification of the most abundant fibril polymorphs from several proteinopathies, including tau, e.g., Alzheimer’s disease (AD)^6^ or Pick’s disease^7^; alpha-synuclein (α-syn), e.g., Parkinson’s disease (PD)^8^, dementia with Lewy bodies (DLB)^8^ and multiple systems atrophy (MSA)^9^; and others. Remarkably, diseases associated with aggregation of the same protein can exhibit different fibril polymorphs. For example, α-synuclein fibrils from patients with PD and DLB adopt the “Lewy fold”^8^, whereas MSA is associated with two different fibril polymorphs (MSA type I and II)^9^.

The finding that the same protein adopts different fibril polymorphs across proteinopathies with distinct pathological signatures suggests a link between structure and function of amyloid fibrils. Therefore, understanding which different polymorphs exist in human brain tissue and where their diversity comes from is essential to solve the link between a fibril’s structure and its potency to spread and cause disease.

Possible causes of polymorph diversity have been proposed, but none have yet been proven. Fibril polymorphs are likely shaped by the cellular environment, as evidenced by structural differences between α-syn fibrils in PD/DLB (predominantly neuronal pathology) and MSA (predominantly oligodendrocyte pathology), as well as *in vitro* studies showing that chemical conditions influence fibril structure^10,11^. Also at the ultrastructural level, aggregates in distinct cells within the same tissue slice display a high level of heterogeneity^12–16^. Distinct polymorphs could potentially even co-exist within the same aggregate, which would explain why differently post-translationally modified α-syn species form layers within a single Lewy body (LB)^17,13^. Currently, structural studies of amyloid fibrils that rely on bulk brain homogenate lack the spatial context needed to clarify the origin of fibril heterogeneity.

Unlike sarkosyl extraction and SPA, the combination of *in situ* cryo-electron tomography (cryo-ET) with subtomogram averaging (STA) preserves spatial context and enables the study of a few fibrils within a single aggregate. However, cryo-ET and STA are technically much more challenging methods in which preserving cellular context generally comes at the cost of lower resolution and a low throughput. Thanks to the development of the serial lift-out technique, cryo-ET can now be applied on whole tissues^18–20^. Thus, although limited in resolution, STA has the potential to connect cellular context with the biochemical and polymorphic diversity of amyloid pathology. This was demonstrated in a recent study presenting the first *in situ* amyloid structures of tau and amyloid-beta (aβ) in postmortem AD brain^21^. Gilbert et al. presented an *in situ* tau fibril structure from Alzheimer’s disease at 8.7Å resolution that was achieved by cryo-electron microscopy of vitreous sections (CEMOVIS^22^), while in the same study, they also used cryo-FIBSEM lift-out, reaching a resolution of 18.1-18.3Å for tau fibril structures.

Given the need to uncover the causes of polymorph diversity and the great pathological heterogeneity in neurodegenerative diseases, *in situ* structural studies in postmortem brain should characterise not only fibril structure, but also the cellular context and biochemical composition of the aggregates they compose. This requires both preserving the tissue ultrastructure and characterising the pathological aggregate, including its cell identity, associated proteins and lipids, and post-translational modifications. Chemical fixation prior to high-pressure freezing (HPF) facilitates this, and is therefore part of the current study.

Here, we present a comprehensive workflow for structural study of *in situ* amyloid fibrils in the postmortem brain, from sample preparation to cryo-correlative light and electron microscopy (cryo-CLEM) and STA. We show that a chemical fixation method which we previously published for a room-temperature CLEM protocol^23^, also preserves tissue ultrastructure after high pressure freezing. By combining immunolabeling with targeted serial lift-out and cryo-focused ion beam (FIB) milling, we used cryo-ET to image tau fibrils in a neurofibrillary tangle (NFT) in an AD brain, as well as α-syn fibrils in a single glial intranuclear inclusion (GNI) in an MSA brain. We further developed a subvolume averaging workflow based on the software Warp, Dynamo^24^ and Relion^25^ to reconstruct tau and α-syn filaments. The resulting densities were compared to existing high-resolution structures solved with SPA. This versatile method enables the study of amyloid fibrils within individual aggregates in fixed postmortem brain tissue and allows connecting cellular and biochemical pathological heterogeneity with structural polymorphism.

## Results

### A cryo-CLEM workflow to specifically target protein aggregates

Preserving the tissue for EM requires minimizing the time between autopsy and fixation, during which the tissue remains susceptible to structural and molecular changes^26^. In this cryo-ET study, we used brain tissue of one AD and one MSA donor (**Extended Data Table 1**) which was chemically fixed using the same protocol as we previously described for room-temperature studies for plastic embedding^23^. In short, after a rapid autopsy protocol (<6h postmortem delay), tissue blocks were dissected in dimensions of 0.5-1 cm in diameter and fixed in 0.1% glutaraldehyde and 4% paraformaldehyde in 0.15M cacodylate buffer (pH 7.4) for 24 hours. The tissue was then stored in 0.1% paraformaldehyde in 0.15M cacodylate buffer (pH 7.4) and shipped from Amsterdam, The Netherlands, to Lausanne, Switzerland, where it was kept at 4°C.

A second main challenge of cryo-CLEM for protein aggregates is the localization and accurate targeting of rare regions of interest, as the final imaged volume is only 20×10×0.2 cubic micrometres. In both the AD and MSA brain tissues, pathology was widespread but precise labeling of these intracellular aggregates for cryo-ET remained essential. After autopsy and fixation, the brain tissue was therefore vibratome sectioned into 40 µm thick slices and immunolabeled with a detergent-free labeling protocol using antibodies targeting pathological post-translational modifications of the aggregated protein species (e.g., phosphorylated α-syn, as shown in **Fig. 1a,b** and **<u>Supp. Video 1</u>**).

**Fig. 1:**
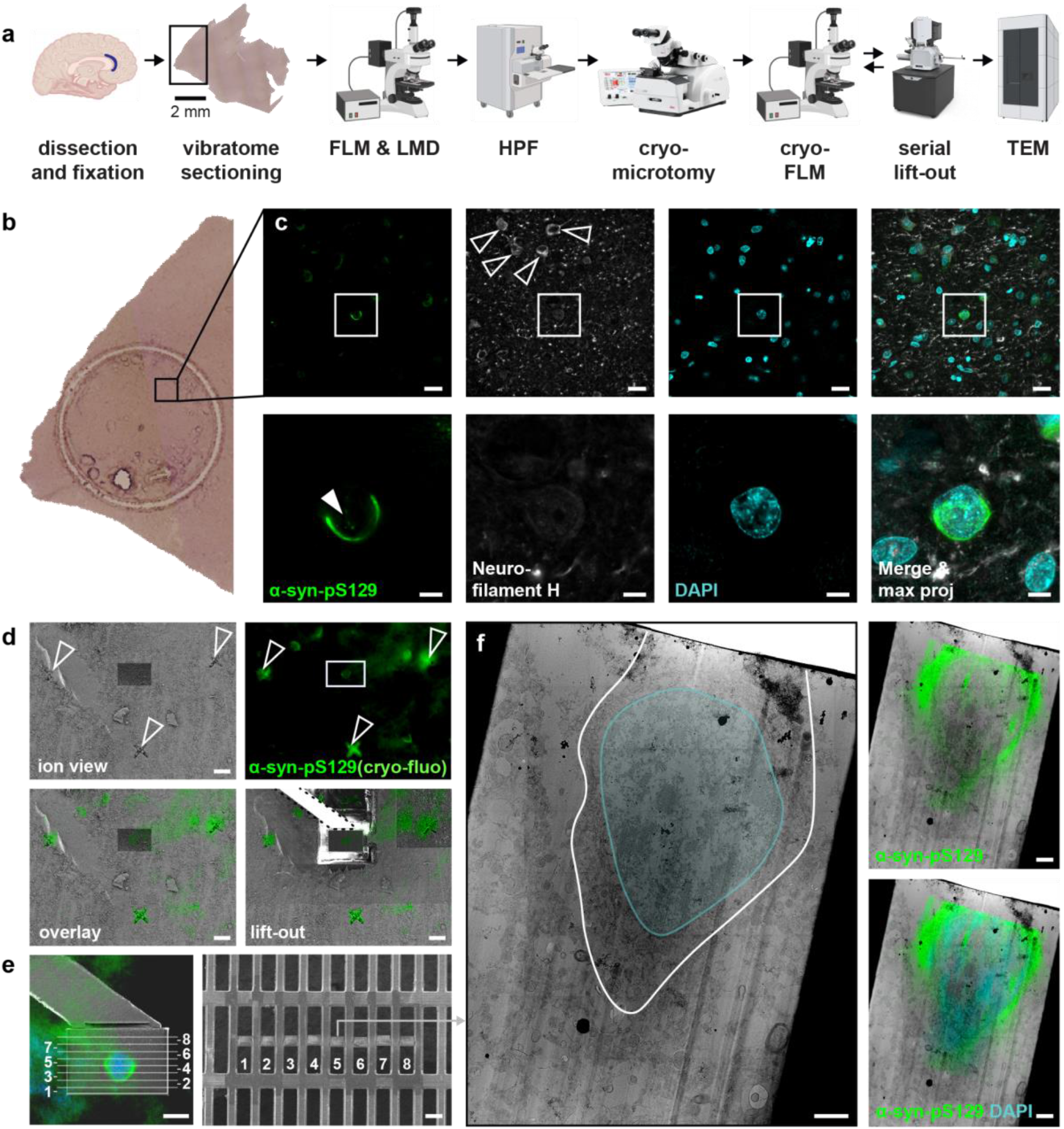
Sample preparation of postmortem human brain tissue for cryo-ET. **(a)** Schematic workflow of the procedure. At autopsy, tissue is fixed in 4% PFA with 0.1% GA for 1 hr. Vibratome sections (40 μm) are immunolabeled and imaged. Disks of 1.5 mm diameter are cut with a laser microdissector. After high-pressure freezing, cryo-microtomy to polish the planchette surface, cryo-fluorescence imaging, and targeted lift-out, lamellae are prepared and imaged in TEM. FLM: fluorescence light microscopy; LMD: laser microdissection; HPF: high-pressure freezing; TEM: transmission electron microscopy. Scale bar: 2mm. **(b)** Close-up of a tissue section of MSA brain tissue after laser microdissection. At the bottom of the disk, a mark made with the LMD serves to recover the disk orientation during high-pressure freezing. **(c)** Fluorescence microscopy image of the glial inclusion that was imaged in TEM. The top row is a large field of view, the bottom row is zoomed in. This glial cell is negative for neurofilament H immunolabeling, as opposed to neurons (open arrowheads). A small intranuclear aggregate is indicated with a closed white arrowhead. Scale bars: top 20 μm, bottom 5 μm. **(d)** Cryo correlation between the ion beam view and the fluorescence image taken with a Leica cryo-Thunder light microscope. First, crosses (open arrowheads) are made surrounding the target cell (white rectangle) with the FIB for alignment, and then a block is prepared for lift-out. Scale bars: 20 μm. **(e)** After lifting the block, eight serial sections with a thickness of 2 μm were made and deposited on a 100-400 rectangular mesh copper grid. Scale bars: left 10 μm, right 50 μm. **(f)** TEM low-magnification search map of the fifth lamella. The cell membrane and nucleus are outlined in white and blue, respectively. Insets on the right show post-correlation with the room-temperature fluorescence image. Scale bar: 2 μm. Schematic illustrations in (a) were made using BioRender.com.

In regions where many aggregates were identified by fluorescence microscopy, disks of ca. 1mm^2^ were cut using laser microdissection and all aggregates within that disk were imaged in 3D with a fluorescence microscope, while still wet and at room temperature (**Fig. 1b,c**). The disk was then high-pressure frozen and re-imaged in a cryo-fluorescence microscope before insertion into a focused ion beam scanning electron microscope (FIBSEM) equipped with a cryo stage. Based on scratch marks on the planchette edge that were visible in both room-temperature and cryo-temperature imaging modalities, cryo-fluorescence images were correlated with the SEM imaging (**Fig. 1d**). For a more precise alignment, cross marks around each aggregate were milled with the ion beam, and the disk was imaged again in cryo - fluorescence to determine the location of each aggregate with respect to the cross marks, following the same procedure as described previously^20^. Serial lift-out of 2 µm thin sections was then performed (**Fig. 1e**). Lamellae were thinned down to ∼200 nm thickness with the FIB, using a 30kV acceleration voltage of the Gallium beam. After transfer of the grid to a 300kV cryo-EM for transmission electron microscopy (TEM), tilt series were taken in regions where filaments were visible. Recorded tilt series images were submitted to alignment and 3D reconstruction using the Warp^27^ and MissAlignment^28^ software.

On low-magnification TEM lamella overviews (searchmaps) where a cellular nucleus was visible, room-temperature fluorescence microscopy images could be post-correlated using the DNA-specific DAPI fluorescence signal and the nuclear envelope visible on the searchmaps (**Fig. 1f**). The good accuracy of this post-correlation confirmed that cryo-CLEM targeting of an immunolabeled aggregate in fixed human brain tissue was possible.

### Chemical fixation and high-pressure freezing preserve tissue ultrastructure

Chemical fixation might introduce crosslinking artifacts, so it is expected that HPF of non-fixed brain tissue is better at preserving proteins in their native state^29^. Immediate HPF directly at brain autopsy is, however, an enormous practical challenge: the technically difficult HPF method has to be performed under bio-safety level 2 (BSL-2) conditions, directly adjacent to the autopsy location. In contrast, fixation in 0.1% glutaraldehyde and 4% PFA, as used in this work, permits a delay between autopsy and HPF while providing the possibility of excellent ultrastructure preservation^23^.

To ensure that the tissue ultrastructure is also preserved when combined with high-pressure freezing rather than EPON embedding, we first inspected the quality of tissue features on the lamellae. Throughout the lifted tissue block, membranes appeared intact, and several features exemplified a good ultrastructural preservation. For instance, nuclear pore complexes (**Extended Data Fig. 1a & <u>Supp. Video 2</u>**), myelin layers (**Extended Data Fig. 1b & <u>Supp.</u> <u>Video 3</u>**), presynaptic vesicles undergoing membrane fusion and postsynaptic densities (**Extended Data Fig. 1c & <u>Supp. Video 4</u>**) were clearly resolved, indicating a good preservation of protein and lipid complexes.

These observations highlight the advantage of chemical fixation at autopsy over freeze-thawing, which could compromise ultrastructure by producing membrane fragments and/or a bursting plasma membrane^21^.

### Tau fibrils in AD brain are consistent with paired helical filaments

Next, we tested whether the structure of the filaments observed within aggregates could be retrieved by subvolume averaging and helical reconstruction (**Fig. 2**). We started with tau tangles in the hippocampus of an AD donor, which are known to contain paired helical filament (PHF) and straight filament (SF) conformations^6,21^. We targeted hippocampal (CA2) tau tangles using an antibody against phosphorylated tau at residue 217 (44-774), which labels pretangles and mature tangles, but not ghost tangles ^16^ (**Fig. 3a,b**). Tau tangles were localized by cryo-light microscopy and imaged by cryo-ET.

**Fig. 2:**
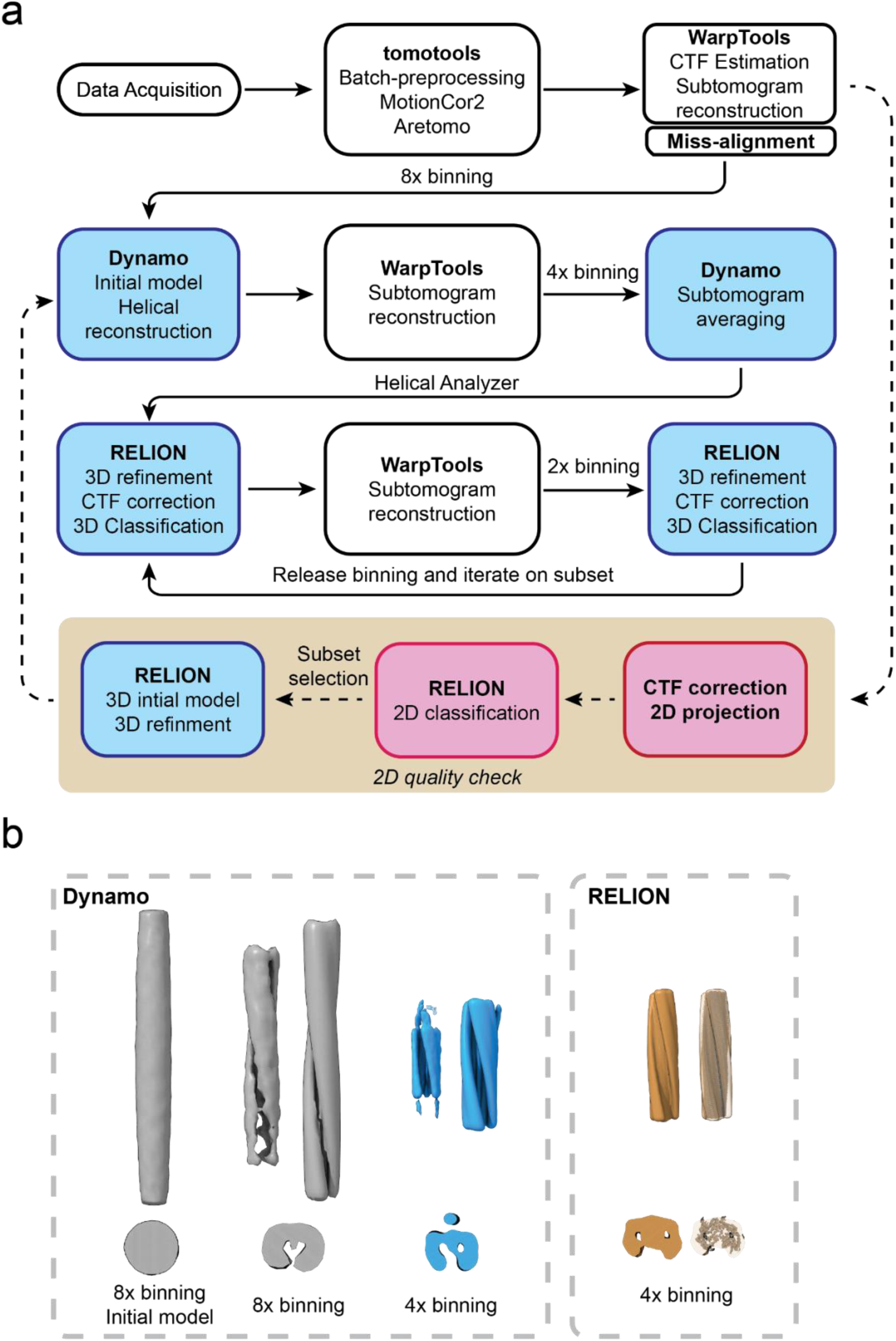
STA workflow for *in situ* amyloid filaments: **(a)** Schematic workflow of STA, indicating the different software packages. Image data were acquired and exported directly from the microscope. The data preprocessing and tomogram reconstruction were done with tomotools and WarpTools, and the tilt-series alignment was improved with MissAlign. The subtomogram alignment and average were done with Dynamo and RELION. Arrows indicate the data flow. **(b)** Initial averages resulting from different data binning stages. STA was first conducted with Dynamo (left panel) with a featureless initial model until binning 4 and then passed to RELION (Right panel). Structures are shown as raw averaged volumes and helical symmetry-applied volumes.

**Fig. 3:**
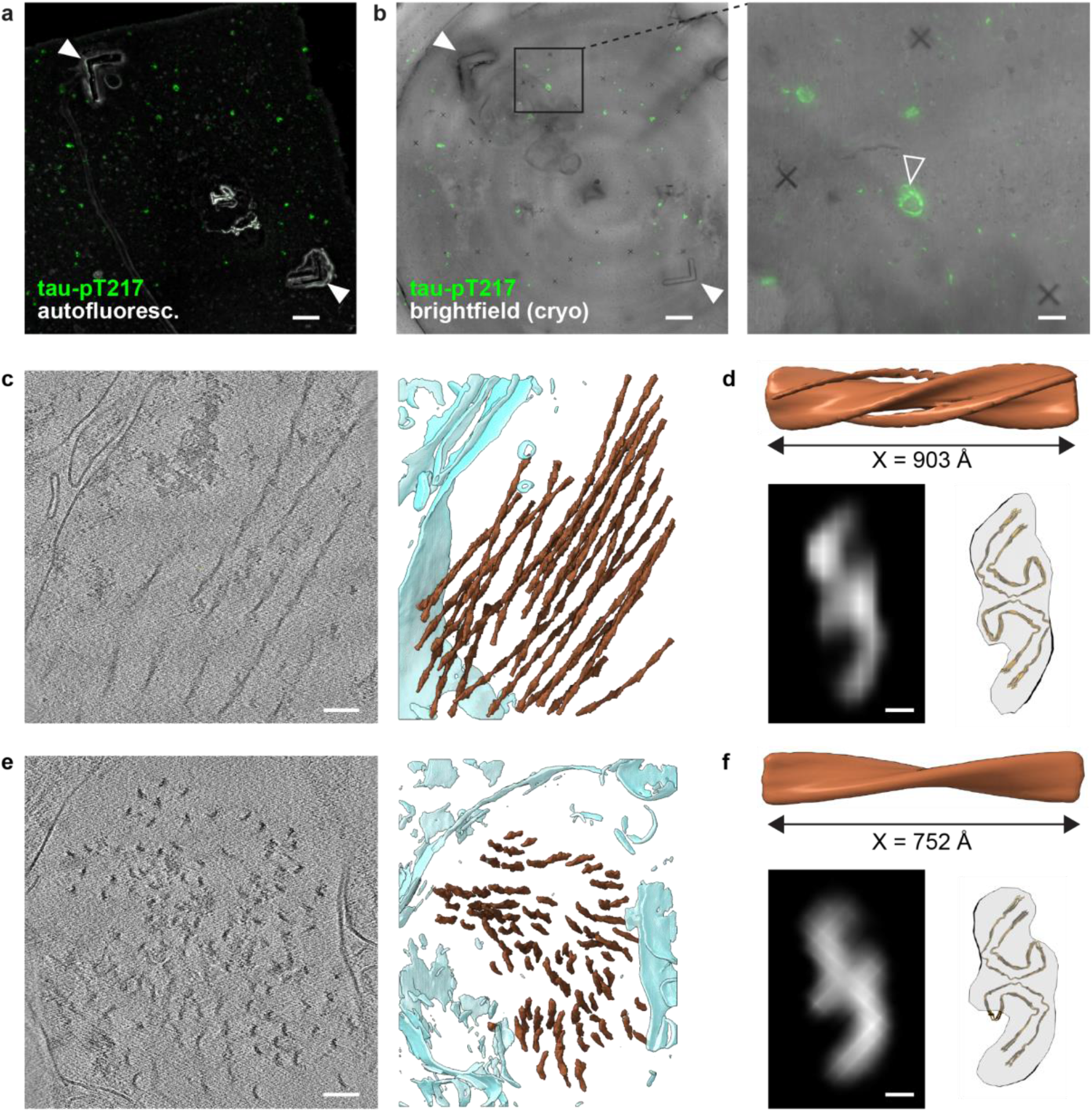
*In situ* tau fibril structures from hippocampal AD brain. **(a)** Fluorescence imaging of tau tangles in the hippocampus. Green: tau-pT217 (antibody 44-744), white: autofluorescence to visualize fiducials made with laser microdissection (white arrows). Scale bar: 100 μm. **(b)** Gray: brightfield image of tissue made in the cryo-thunder after high-pressure freezing, showing that the same fiducials are visible (white arrows). These fiducials are used for post-correlation of the tau-pT217 signal (green). The inset (right) shows the tau tangle targeted for lift-out (open arrow) in between three fiducial crosses made with the FIB. Scale bars: left 100 μm, right 50 μm. **(c)** Left: x-y slice through a reconstructed tomogram; Right: rendering of its segmentation. Scale bar: 100 nm. **(d)** STA reconstructed tau fibrils in the tomogram shown in (c), with the structure of tau PHFs (PDB-5O3L) overlaid for comparison. Scale bar: 20 Å. **(e)** X-y slice through another tomogram on a different lamella (left) with its segmentation (right). Scale bar: 100 nm. **(f)** STA reconstructed tau fibrils in the tomogram shown in (e), again overlaid with PDB-5O3L. Scale bar: 20 Å. Colors in (c) and (e): tau filament = brown, cell membrane = turquoise.

On the tomographic reconstructions, tau fibrils were segmented in IMOD^30^ and subjected to a subvolume averaging procedure. Briefly, this workflow consisted of the following steps (**Fig. 2a**): (1) extraction of subtomograms at a pixel binning of 8 (pixel size 16-20Å) with WarpTools and an initial particle alignment in Dynamo^24^, (4) re-extraction of aligned particles at a binning of 4 (pixel size 8-10Å) and another alignment in Dynamo, (5) removal of particles with unreasonable helical parameters with home-developed software, Rohlex (**Extended Data Fig. 2**), and (6) particle refinement in Relion^31^. For a more detailed overview of the workflow, including the exact processing commands, see the supplementary information and the tutorial at https://lbem-ch.github.io/amy-subtomo-averaging.

To register any potential diversity of fibrils in different aggregates, only fibrils visible on the same tomogram were averaged, and we never computed an average of fibrils extracted from different tomograms. From one tomogram, the density obtained after initial alignment in Dynamo at a binning of 8 (pixel size 19.4Å) accommodated the PHF model (PDB-5O3L). Further alignment at a binning of 4 (pixel size 9.7Å) confirmed that these fibrils were PHFs, with a final density map at 35Å resolution (**Fig. 3c,d & <u>Supp. Video 6</u>**). Of note, re-extraction of the particles at a lower binning and subsequent averaging was tried extensively but did not yield any higher resolution averages.

Another tomogram on a different lamella originating from the same lifted block displayed tau fibrils perpendicular to the lamella plane, which appeared to be in a neurite (**Fig. 3e,f & <u>Supp.</u> <u>Video 7</u>**). The resulting density again was reminiscent of the symmetric “double C” of the PHF. For both tomograms analysed, 3D classification did not separate any other classes, suggesting that all tau fibrils averaged are of the PHF type rather than SF.

In addition to tau filaments in the AD brain, we validated the subvolume averaging workflow on cytoskeletal filaments in the postmortem brain of an MSA donor, which were not targeted but found adjacent to aggregates in the cingulate gyrus. Actin filaments in a dendritic spine were found to adopt the expected twist and rise after an initial alignment in Dynamo at a binning of 8 (pixel size 15.84Å), and the density eventually reached a resolution of 30 Å (**Extended Data Fig. 1c-e**). In addition, processing of neurofilament bundles in a neurite resulted in a density resembling a pentamer (**Extended Data Fig. 2f-h**), similar to vimentin intermediate filaments^32^. This similarity motivated us to apply the same helical symmetry to neurofilament as to vimentin, resulting in a density with a resolution of 36 Å (**Extended Data Fig. 2i & <u>Supp. Video 5</u>)**.

Together, these results demonstrate that subvolume averaging can recover disease-relevant fibril architectures directly from intracellular aggregates *in situ*. In the analysed AD tau tangles, the reconstructed densities were consistent with the PHF fold. The reconstruction of actin and neurofilament assemblies further demonstrates that this approach is not restricted to tau, but can be broadly applied to structurally characterise helical filaments in postmortem human brain tissue.

### α-Syn fibrils in a glial nuclear inclusion best fit the MSA type-I polymorph

Next, we applied the same procedure to study α-syn fibrils in the cingulate gyrus of an MSA donor. Aggregates rich in α-syn phosphorylated at residue 129 (pS129) were stained with the EP1536Y antibody, as showcased in **Fig 1**. The brain of the same patient was previously characterised by us using room-temperature CLEM encompassing EPON embedding, which confirmed the presence of neuronal and glial aggregates rich in α-syn fibrils^33^. Here, we aimed to specifically target aggregates in oligodendrocytes (GCIs and glial nuclear inclusions (GNI)), which are thought to cause demyelination and subsequent neurodegeneration in MSA. Because neuronal pathology is also present in MSA brain^34^, although less abundant than GCIs, we co-labeled the tissue with an antibody targeting neurofilament. All flame-like aggregates that were negative for the neurofilament marker were considered to be GCIs and targeted for cryo-FIBSEM lift-out after high-pressure freezing.

By immunofluorescence, we found that most oligodendrocytes with cytoplasmic aggregates also contained phosphorylated α-syn in the nucleus, where it formed small aggregates of less than 1 µm diameter (white arrow in **Fig. 1c**). It is not known how these GNIs develop and whether they are structurally similar to their cytoplasmic counterparts.

The nuclei of oligodendrocytes could clearly be identified on low-magnification cryo-TEM overviews (“search maps”). In most regions with high pS129 signal, fibrils with a thickness of about 20 Å could be identified, both surrounding the nucleus (as GCIs) and within the nuclear envelope (as GNIs). As opposed to tau PHFs, the twist of α-syn fibrils was not directly apparent even from high-magnification cryo-EM micrographs. We thus proceeded by recording tilt-series at a pixel size of 1.98 Å and a total dose of 153 e/Å^2^.

We selected a tomogram of a GNI for further processing, with fibrils oriented perpendicular to the lamella plane (**Fig. 4a,b & <u>Supp. Video 8</u>**). An initial subvolume alignment in Dynamo of particles extracted at a binning of 8 yielded a fibrillar density with a major groove displaying a left-handed twist of –1.4 degrees at an estimated rise of 4.8Å, which is in accordance with the twist and rise of α-syn fibrils extracted from the MSA brain^9^.

**Fig. 4:**
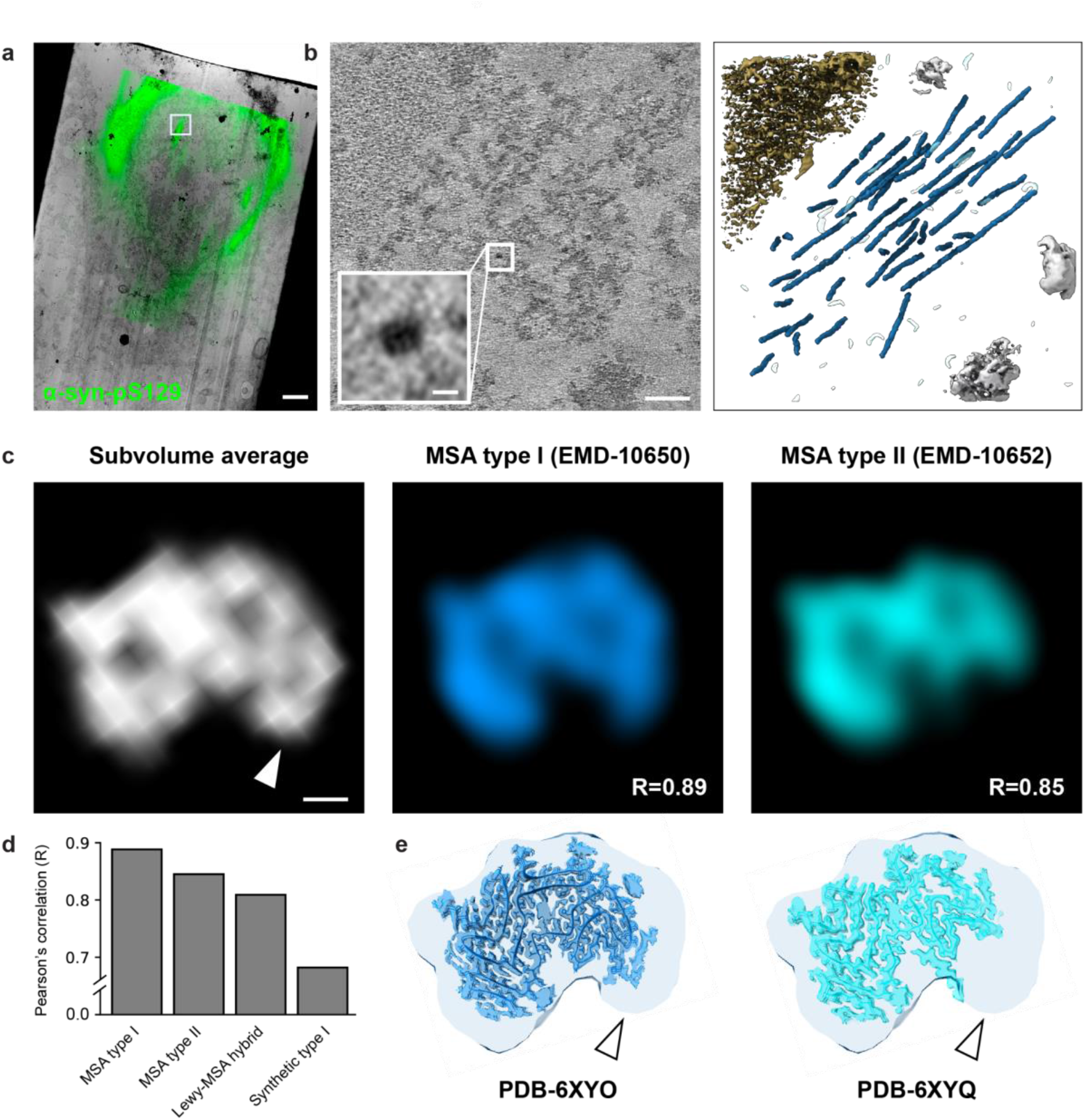
*In situ* α-syn subvolume average of α-synuclein fibrils in a single glial intranuclear inclusion is best accommodated by the MSA type I fibril. **(a)** Low-magnification TEM micrograph post-correlated with fluorescence microscopy (post-correlation was made based on the DAPI signal). The white square indicates the position of the tilt-series acquisition. Green: α-syn-pS129 (antibody EP1536Y). Scale bar: 2 μm. **(b)** Left: Tomographic slice showing fibrils perpendicular to the imaging plane. The inset shows an enlarged cross-section of a fibril. Right: tomogram segmentation with fibrils (blue), nucleosome-like structure (yellow) and aggregates in gray. Scale bars: 100 nm, inset: 10 nm. **(c)** Top view of the subvolume average of intranuclear fibrils from the tomogram in (b), next to the top views of densities of *ex vivo* fibrils from the MSA brain filtered to a resolution of 30 Å (MSA types I and II, EMD-10650 and 10652 respectively ^9^). Scale bar: 20 Å. **(d**) Barplot comparing the Pearson’s correlation coefficients of filtered densities fitted in the subvolume average. **(e)** Top views of the subvolume density, overlaid with MSA types I and II non-filtered densities and atomic models. White arrows indicate the flanking density better explained by MSA type I than type II.

When we re-extracted these particles at a binning of 4 and re-aligned them in Dynamo, we noticed that a large proportion of these particles (∼44%) had refined Euler angles that were not in accordance with neighboring particles within the same fibril (**Extended Data Fig. 2**). In some cases, an entire fibril consisted of particles that had irregular rotation angles, which we interpreted as a sign of the absence of a regular twist. Whether those fibrils were indeed non-twisting, or whether the signal-to-noise ratio is not high enough to determine their twist, remains to be determined. Nonetheless, we deleted those poorly aligned subvolumes from the analysis using a home-developed software called Rohlex, leaving only 5312 subvolumes (out of the original 9422) for subsequent refinement in Relion (**Extended Data Table 2**).

The final density of fibrils in this GNI reached a resolution of 30Å and accommodated the MSA type I and II filament structures (PDB-6XYO and 6XYQ, respectively). To estimate the accuracy of the fit, we lowpass filtered the electron densities corresponding to these atomic models (EMD-10650 and 10652) to the same 30Å resolution of the STA density (**Fig. 4c**). We then fitted the lowpass filtered maps to the STA density using ChimeraX, which minimises the cross-correlation, and used the Pearon’s correlation coefficient (R) of voxels within a tight mask (with a height of 30% of the crossover distance) as a measure of how well the known MSA structures are accommodated by the STA density.

We found the R of lowpass-filtered MSA type I and II densities to be 0.89 and 0.85, respectively. For comparison, the recently published “Lewy-MSA hybrid fold” (EMD-70295), which was extracted from an atypical MSA patient with a high amount of neuronal aggregates^35^, yielded a correlation of 0.81, and a double protofilament synthetic α-syn fibril of type 1 (EMD-19986) yielded a correlation of 0.68. The latter two structures both look visually very different from the STA density (**Extended Data Fig. 3**). Of note, 3D classification of the particles did not identify multiple fibril types within this tomogram, and we suspect all fibrils in this aggregate to be of the same type.

Both by visual inspection and measured by R, the MSA type I fibril best explains the density obtained by STA. However, the resolution obtained is too low to unambiguously attribute either the type I or type II filaments to the GNI fibrils. Of note, one specific region in the density was particularly poorly explained by the MSA type II fibril (white arrows in **Fig. 4c**), but it could accommodate the density flanking the C-terminal end of the type IA protofilament, which was not modeled in PDB-6XYO. With the current data, however, it remains impossible to determine the exact identity of this density. Future studies comparing fibrils across multiple GNIs and GCIs will be needed to validate this observation and determine the distribution of type I and type II fibrils within these inclusions.

### Preserved ultrastructure and STA reveal α-syn fibrils crossing the nuclear envelope

Next to GCIs and GNIs alone, one observation deserving attention is that of an α-syn fibril penetrating the nuclear envelope. This fibril was found in a glial cell different from that shown in **Fig. 4**, which also contained a perinuclear GCI and a small (<1μm) GNI (**Fig. 5a**). The DAPI signal revealed an unusual nuclear shape, with a nuclear invagination disturbing its otherwise elliptical morphology (white arrow in **Fig. 5a**). At the edge of the nucleus, TEM revealed a ∼20Å-diameter fibril oriented almost parallel to the lamella traversing the nuclear membrane, which folded outwards into the cytoplasm and appeared highly disrupted (**Fig. 5b-c & <u>Supp.</u> <u>Video 9</u>**). The enriched protein density surrounding the fibril at the crossing point and the disrupted nuclear envelope prevented us from determining whether an NPC was present.

**Fig. 5:**
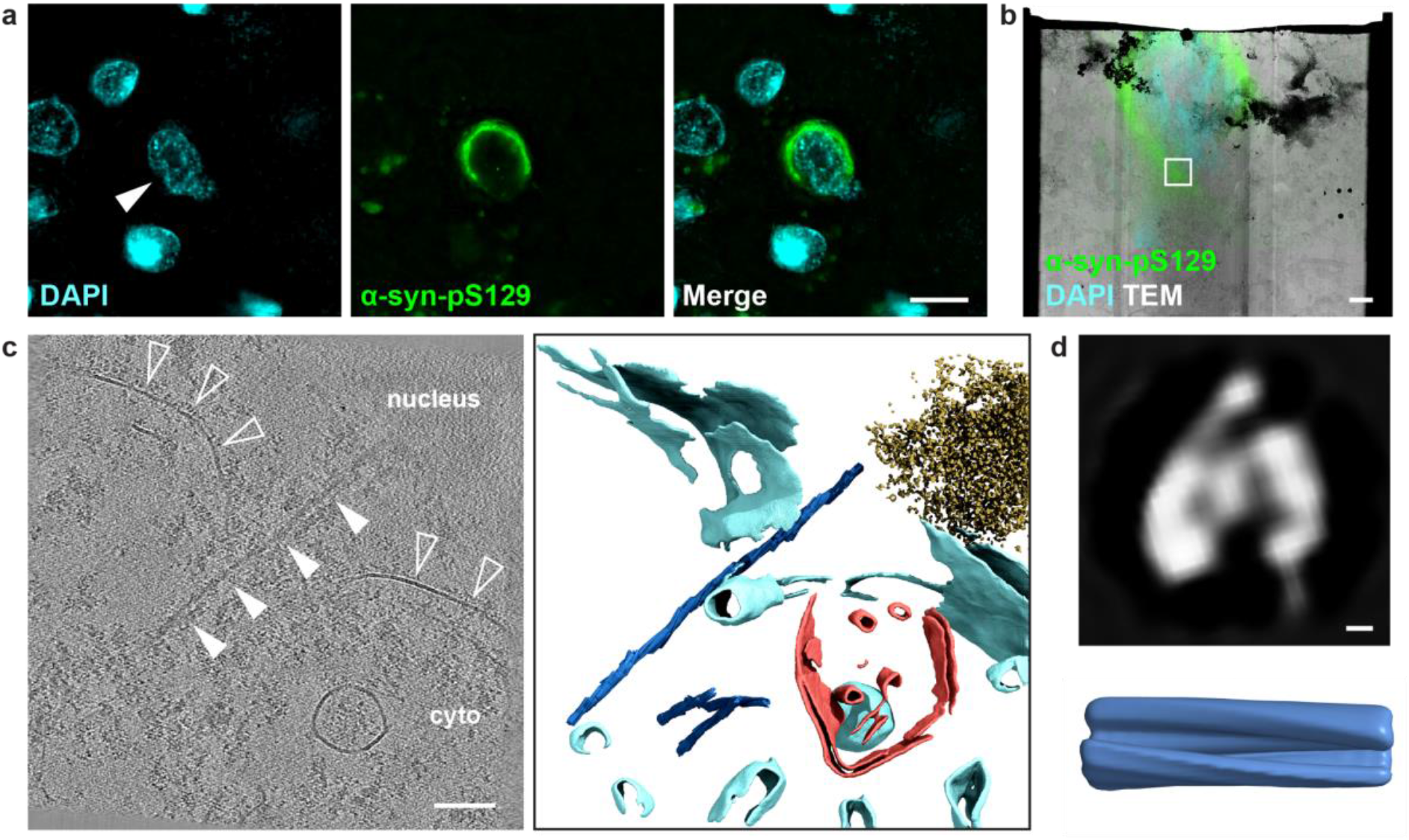
An α-syn fibril penetrates the nuclear envelope. **(a)** Fluorescence microscopy of the glial cytoplasmic inclusion targeted for serial lift-out. The white arrow indicates a small indentation of the nuclear shape. Scale bar: 10 μm. **(b)** Post-correlation of the immunofluorescence signal (cyan: DAPI, green: α-syn-pS129) and the low-magnification TEM overview (searchmap). The white square indicates the position of the tilt-series acquisition. Scale bar: 2 μm. **(c)** Left: Tomographic slice showing a fibril (closed white arrows) penetrating the disrupted membrane of the nuclear envelope (open arrows). Right: segmentation showing fibrils (dark blue), membranes (light blue), nucleosome-like structures (yellow) and mitochondrial membranes (red). Scale bar: 100 nm. **(d)** Top view of the subvolume average of the same fibrils shown dark-blue in (c), with the side-view shown below. This average was made in Dynamo and, due to a too low particle number, could not be refined in Relion. Nevertheless, its shape is similar to the intranuclear fibrils after Dynamo alignment (Fig. 2b).

To confirm the identity of the fibril, we segmented it together with three similarly sized fibrils perpendicular to the lamella and subjected all four to subvolume averaging. Due to a low particle number (1584) and a poor signal-to-noise ratio, gold-standard refinement in Relion did not yield any interpretable density, and only the Dynamo alignment at a binning of 4 (pixel size 7.92Å) could be used for comparison. The resulting density (**Fig. 5d**) displayed the same “3”-shape observed for the α-syn fibrils in **Fig. 4**, confirming that the fibril penetrating the nuclear envelope is α-syn. Thus, these data provide direct ultrastructural evidence of α-syn fibrils traversing the nuclear envelope, highlighting a potential route by which fibrillar α-syn could access the nucleus.

Overall, cryo-ET of chemically fixed and immunolabeled brain tissue enabled us to identify PHFs in AD tau tangles and to attribute MSA fibrils in a GNI to most likely MSA type I filaments. Although the low-resolution densities remain difficult to interpret, reminding us of the era of “blobology”, the availability of high-resolution amyloid structures solved by SPA provides structural references for assigning STA densities to the most likely polymorphs. Importantly, preservation of the native ultrastructure also enabled observations that would otherwise be impossible, such as an α-syn fibril penetrating the nuclear envelope, providing mechanistic insight into how GNIs may form.

## Discussion

Previous cryo-EM studies reported that amyloid fibrils adopt disease-specific polymorphs, and that these structures are the same in different individuals with the same disease. However, a certain degree of polymorph heterogeneity within a single patient has also been reported. *In situ* cryo-ET has the potential to unravel the biological origin of this polymorphism within a single patient, if it is combined with a biochemical and spatial characterization of the fibrillar aggregate studied by STA. The workflow we present here makes that possible by immunolabeling and fluorescence imaging of human brain tissue prior to HPF.

Multiple biological questions can be addressed because, in many cases, fibrils differ sufficiently in morphology to be distinguished by cryo-ET in the fixed brain, despite the lower resolution. For example, PHFs versus SFs are distinguishable at low resolution and were both identified in the hippocampus of the AD brain^21^. Our study only identified PHFs, which could indicate an increased labeling efficiency for PHFs of the tau antibody we used to target NFTs (44-774). Broader analysis will be required to establish the prevalence of PHF and SF conformations across aggregates and relevance of the two distinct fibril types for AD. For α-syn, our density supports the MSA type I filament as the most likely candidate for the fibrils within the studied GNI. Other MSA-relevant folds, such as the Lewy-MSA hybrid, showed worse agreement with the density. Whether this polymorph is indeed characteristic of neuronal pathology as proposed by Enomoto et al.^35^ could be tested by expanding our cryo-ET workflow to neuronal aggregates in a larger cohort of MSA patients. Further, cryo-ET applied to fixed PD brains could help us understand the structural differences (if any) of fibrils in LBs, pale bodies and Lewy neurites, as well as the spatial distribution of twisting versus non-twisting fibrils in the brain. Finally, our approach could help investigate the pathological significance of single versus double protofilaments, which have been shown to co-exist in PD patients bearing the A53T mutation^36^ and also in mouse models of synucleinopathies^37^.

Compared to other methods providing ultrastructural characterization, particularly room-temperature CLEM of heavy-metal-stained and EPON-embedded tissue^23^, cryo-ET has the major advantage of allowing the identity of observed fibrils to be established. After heavy-metal staining, some filaments are morphologically too similar to distinguish. For example, α-syn fibrils and neurofilaments have roughly the same diameter and often occur in close proximity^13,15^. This makes them difficult to distinguish and limits the interpretation of room-temperature CLEM. As our data shows, cryo-ET combined with subvolume averaging can distinguish α-syn fibrils from neurofilaments. This makes it possible to confirm the identity of filaments in new observations. As an example, we presented a tomogram of an α-syn fibril traversing the disrupted nuclear envelope. A similar observation could have been made in EPON-embedded tissue, but the identity and polymorph of the filament could not have been determined. Here, this observation may provide a morphological basis for the formation of nuclear inclusions from cytoplasmic α-syn fibrils. This is particularly relevant given the absence of a mechanistic model for GNI formation in MSA.

The main limitation of the presented workflow remains the low resolution of the densities obtained. The 35Å tau PHF density presented here is at much lower resolution than the 8.7Å reported by Gilbert et al., which came from unfixed brain tissue cryo-sectioned by CEMOVIS^21^. Aldehyde fixation, although facilitating immunolabeling and long-term storage of the tissue at 4 degrees, may contribute to the low resolution. Other factors beyond fixation may also contribute to this difference, including fibril heterogeneity, differences in lamella preparation (CEMOVIS versus FIBSEM), specifics of the TEM tilt-series acquisition, or prolonged tissue storage in buffer (2-3 years). Indeed, crosslinking by fixation may not substantially alter amyloid fibril structure, as brain extracts from MSA patients retain prion-like seeding activity following aldehyde fixation^38^. The latter work suggests that even though chemically fixed human brain is classified as BSL-1, care should be taken when handling that material.

Overall, while the origin of fibril polymorphism in the brain of single individuals remains elusive, *in situ* cryo-ET with sub-volume analysis of amyloids can help to study it. To do so, aldehyde fixation and immunolabeling enable targeting intracellular aggregates with a defined composition. Careful inspection of the aligned Euler angles of the subvolumes and removal of the misaligned particles is necessary to obtain a reliable density which, even if the resolution is low, can be put in context with the already existing high-resolution structures obtained through sarkosyl extraction and SPA. Extending this method to a diverse cohort of brain donors and neurodegenerative diseases will enable a deeper understanding of the biochemical and structural heterogeneity of amyloid pathology.

## Methods

### Postmortem human brain samples

Postmortem human brain tissue was collected from donors enrolled in the Netherlands Brain Bank donation program (www.brainbank.nl) using a rapid autopsy protocol (<6h postmortem delay). All donors had provided written informed consent for brain autopsy and the use of their tissue and clinical data for research. All procedures were performed in accordance with the declaration of Helsinki. Detailed neuropathological diagnoses and clinical information were obtained following local ethical and legal guidelines, and all procedures were approved by the Amsterdam UMC institutional review board (NBB 2019/148). The use of human post-mortem tissue at EPFL (Lausanne, Switzerland) was approved by the Cantonal Ethics Committee of Vaud (CER-VD; approval numbers 2020-01692 and 2025-01061), in accordance with Swiss Human Research Act guidelines. We included hippocampus (CA2) tissue from a pathologically confirmed AD and anterior cingulate cortex from a MSA case (see **Extended Data Table 1** for details).

### Immunolabeling and high-pressure freezing

Directly at autopsy, tissue blocks of 0.5 to 1.0 cm diameter were dissected from regions of interests and fixed in 0.1% glutaraldehyde and 4% paraformaldehyde in 0.15M cacodylate buffer for 24 hours and stored long-term in 0.15M cacodylate buffer with 0.1% paraformaldehyde. With a vibratome (Leica VT1200), 40 μm thick free-floating sections were made. Hippocampal tissue from the AD brain donor was immunolabeled with a primary antibody targeting tau-pT217 (44744 - Thermo Fisher Scientific) and AmyTracker 680 (Ebba Biotech). Cingulate gyrus tissue from the MSA brain donor was immunolabeled with primary antibodies against α-syn-pS129 (EP1536Y - Abcam # 51253 - 1/1000) and neurofilament (Sigma Ab5539 - 1/500). All primary antibodies were diluted in TRIS-buffered saline (TBS) and the tissue slices were incubated at 4°C overnight. After three washes in TBS, the tissue was stained for 1 hour with Alexa-conjugated secondary antibodies (donkey 488 Molecular Probes A32787 - 1/400, chicken 647 Molecular Probes A32933 - 1/400) and DAPI (Biolegend #422801 - 1/800 dilution) at room temperature. Tissue was mounted on a glass slide embedded in medium of glycerol/TBS (1:1; pH 7.4) with a coverslip on top, and fluorescence microscopy images were taken with a Thunder Tissue imager (Leica Microsystems) using a 63 x oil objective (HC PL APO/1.4 numerical aperture).

Sections were then demounted from the glass slide and mounted on a ca. 400 nm thick polyetherimide (PEI) film, which was custom made on a metal Leica film support. A second film was mounted on top, so that the tissue was sandwiched between the two films. In regions with abundant pathology, correlation marks were made with a laser microdissector (Leica LMD7), and disks of ca. 1mm^2^ were sectioned and collected in TBS buffer with 20% dextran in which they were stored overnight at 4°C. The disks were transferred in a 50 µm deep cavity of a 3 mm aluminum carrier (Art.390, Wohlwend GmbH, Switzerland) filled with 20% dextran in TBS. The carrier was covered with a tap carrier (Art.1323, Wohlwend GmbH, Switzerland) and vitrified in a Leica EM ICE high-pressure freezer (Leica Microsystem, Vienna, Austria). After freezing, carriers were stored in a cryobox in liquid nitrogen until further processing.

### Cryo-CLEM and Serial Lift-Out

The full cryo-CLEM procedure was done as described earlier^20^. Briefly, after high-pressure freezing, the surface of the frozen sample is too rough for cryo-CLEM and first needs to be polished. To do so, the carrier was loaded into a Leica UC6 cryo-ultramicrotome (Leica Microsystems, Vienna, Austria) cooled down to –190°C (chamber) and gently trimmed using a Trim20 diamond knife (Diatome, Biel, Switzerland). For correlation, marks were engraved on the carrier rim using a diamond tip. The carrier was then mounted into a Leica cryo-Thunder light microscope (Leica Microsystems, Vienna) to re-image the protein aggregates with a 50X objective (0.75 numerical aperture) and take an overview of the full carrier and rim. It was next loaded into the Aquilos 2 cryo-FIBSEM (Thermo Fisher Scientific Inc., US), where it was first sputter-coated with platinum at 1 kV, 30mA, 0.15 mbar for 15 seconds to reduce charging effects. Using the ion beam, an overview montage was then made and aligned with the cryo-thunder light microscope image using the MAPS software (Thermo Fisher Scientific Inc., US), to establish roughly the positions of the protein aggregates. With the FIB, three cross-shaped fiducial markers were then milled surrounding each aggregate with a 1nA ion beam current, and the carrier was transferred back to the Leica cryo-Thunder to re-image the pathology again including the fiducial markers. It was then re-loaded into the Aquilos 2 cryo-FIBSEM in position 1 of the shuttle, together with a copper receiver grid (100-400 rectangular mesh, Agar AGG2140C) in position 2. A new ion beam overview montage was made and aligned to the images made in the second cryo-Thunder acquisition. This alignment was very precise thanks to the cross-shaped fiducials. A block of 40 x 20 μm, with the aggregate of interest in its center, was prepared with a milling procedure as described^20^. It was then attached to a silver needle, lifted, and a layer of organometallic platinum using the gas injection system (GIS) was applied by opening the flow for 4×20 seconds. The block width was then adapted to fit the grid bar spacing and was transferred to the receiver grid, where lamellae of ca. 2 μm thickness were made by first attaching the bottom of the grid to the sides of the grid bars, cutting the lamella, and gently lifting the block. Typically, about 8 lamellae could be prepared per block.

The lamellae were thinned with the FIBSEM Gallium^2+^ beam at an acceleration voltage of 30kV, using the AutoTEM software (Thermo Fisher Scientific Inc., US). The following three milling currents were applied using automated AutoTEM milling: 1nA to reach a thickness of 2μm, 0.5 nA to reach 1.4μm and 0.3nA to reach 800 nm. The final polishing was done manually at 0.1nA and 50pA using rectangular milling patterns and an overtilt of 0.7° to thin the far side of the lamella.

### Cryo-Electron Tomography

The grid was loaded into the Titan Krios G4 equipped with a Selectris X energy filter and a Falcon4i direct electron detector camera (Thermo Fisher Scientific Inc., US). Image acquisition was done using the EPU software (Thermo Fisher Scientific Inc., US). First, overview montages of the entire lamella were acquired at 15’000x magnification (pixel size 22.4Å). In regions with fibrils, tilt series were acquired at 53’000x magnification for tau (pixel size 2.42Å) and 64’000x magnification for α-syn (pixel size 1.98Å), a 50μm C2 aperture and an energy slit width of 10eV. A dose-symmetric tilt scheme was used with 3° increments, with a dose of ∼3.8 e^-^/Å^2^ per tilt and a target defocus range of –1,…,–5 μm. The tilt range spanned from –50° to 70°, resulting in a total dose of 153-155 e^-^/Å^2^. See **Extended Data Table 2** for the acquisition specifics of each analysed tomogram.

Tilt-series were motion-corrected with Tomotools^39^, aligned with AreTomo^40^ and MissAlignment^28^, and reconstructed with WarpTools^27^, as detailed in the Supplementary Information. Membrane segmentations were made with MemBrain-v2^41^ and nucleosome segmentations by low-pass filtering and thresholding the raw tomogram. Filaments were rendered onto the segmentations using ArtiaX^42^ in ChimeraX^43^.

### Subtomogram averaging for *in situ* filaments in Cryo-ET

A step-by-step description of the subvolume averaging pipeline can be found in the

## Supplementary Information

### Helical filament analyzer

3D classification for *in situ* STA is difficult due to the lower S/N ratio. To analyze and classify helical filaments, we developed a software tool (Rohlex for “rolling helix”) to visualize and modify subtomogram alignment results from RELION and Dynamo. For each filament segment, its coordinates and pose are back-mapped into the tomogram to determine its position and azimuthal angle along the filament axis. A model helix defined by the user-specified helical rise and twist is then plotted as a reference to evaluate whether the aligned segments follow the expected helical arrangement.

In brief, the axis of each filament (V) is determined from the spatial distribution of its segment coordinates, with the mean coordinate defining the center of the filament. Since each segment is refined by aligning a reference whose z-axis lies along the filament, its orientation defines a segment direction (Vi) and an azimuthal angle (Ri) around V. The relationship between V and Vi indicates whether a segment points along, opposite to, or away from the filament axis, while Ri describes how the segments rotate along V. Ideally, the measured orientations follow the helix defined by the user-specified rise and twist. By comparing the aligned segments with this model reference, users can generate exclusion and pose-correction lists for subsequent RELION or Dynamo alignment jobs.

For easier illustration, we wrapped the Ri angle in Fig. S2 (Y-axis). Detailed methods and implementation are available on GitHub: https://github.com/LBEM-CH/rohlex

## Data availability

Raw cryo-ET tilt series will be made available on the EMPIAR database upon acceptance of the manuscript. The 3D density maps will be made available on the EMDB upon acceptance of the manuscript. A tutorial of the full tomogram reconstruction and subtomogram averaging workflow is available at https://lbem-ch.github.io/amy-subtomo-averaging. All scripts are available from the github repository related to this tutorial: https://github.com/LBEM-CH/amy-subtomo-averaging. The Rholex software is available at https://github.com/LBEM-CH/rohlex. The BioRender Licence number will be added upon acceptance of the manuscript.

## Author contributions

L.v.d.H., W.-L.C., A.-L.M.-M. and H.S. designed the study. W.D.J.v.d.B. performed rapid autopsies of brain donors, collected brain tissue for EM and performed neuropathological assessment. L.v.d.H., D.A.S., N.S. and A.J.L. immunolabeled brain tissue and performed fluorescence light microscopy. J.D. performed cryo-FIBSEM lift-out with C.G., L.v.d.H. and D.A.S. providing assistance. J.R., L.v.d.H., D.A.S and W.-L.C. collected cryo-TEM tilt series. W.-L.C. developed the software and data analysis pipeline. W.-L.C., L.v.d.H. performed data analysis. L.v.d.H., W.-L.C. and H.S. wrote the manuscript. All authors contributed to the analysis and interpretation of the data and approved the final version of the manuscript.

## Acknowledgements

We are grateful to the individuals who participated in the brain donation program of the Netherlands Brain Bank and their families, making this study possible. We thank the staff at the EM Facility at the University of Lausanne (UNIL), as well as the staff of the Dubochet Center of Imaging Lausanne for support in sample preparation and image data collection. We thank Babatunde Ekundayo (EPFL) for his encouraging support. We thank Gaël Cartier-Michaud from the EPFL for expert IT support.

## Funding

This work was supported by the Swiss National Science Foundation, grant 200021_200628 and by the European Union (ERC 4D-BioSTEM, No. 101118656). Views and opinions expressed are, however, those of the authors only and do not necessarily reflect those of the European Union or the European Research Council Executive Agency. Neither the European Union nor the granting authority can be held responsible for them.

## Competing interests

The authors declare no competing interests.

## Extended data

**Extended Data Table 1:** Demographic and neuropathological details of brain donors in this study.

| Donor ID | A | B |
| --- | --- | --- |
| Clinical / pathological diagnosis | MSA-P / MSA-P | AD / AD |
| Brain region | CG | HIP/CA2 |
| F/M | M | M |
| Age | 68 | 70 |
| Postmortem delay | 4h45m | 4h50m |
| ABC score<sup>44</sup> | A1B1C0 | A3B3C3 |
| Braak stage (α-syn) | 6 | 0 |
| Braak stage (tau) | 0 | 6 |
| Time kept in storage buffer | ~4y | ~1y |

MSA=multiple systems atrophy, AD=Alzheimer's disease, CG=cingulate gyrus, HIP=hippocampus, CA2=cornu ammonis 2, F=female, M=male.

**Extended Data Fig. 1:**
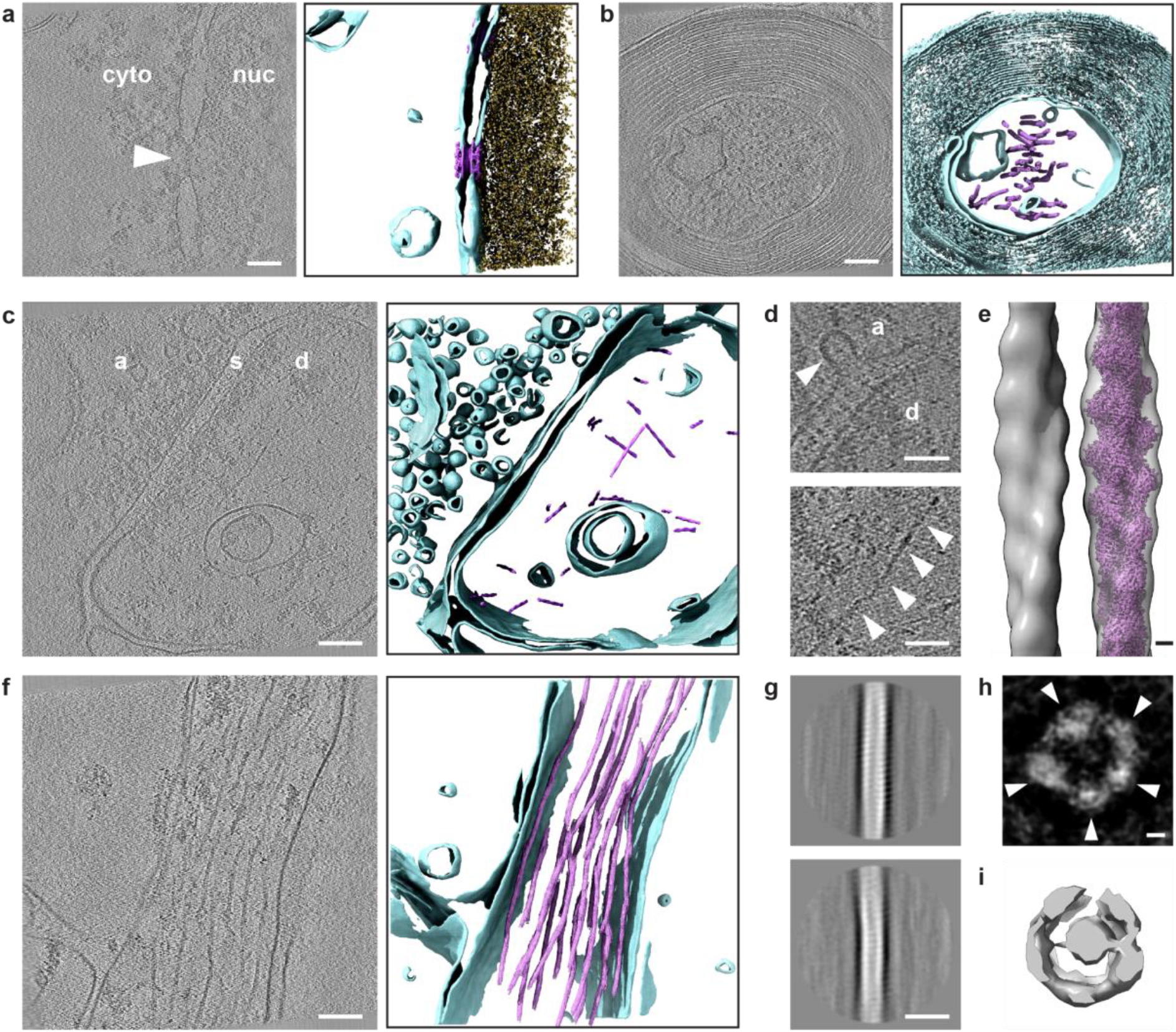
Ultrastructural preservation of chemically fixed and high-pressure frozen postmortem MSA brain tissue. **(a)** Tomographic slice (left) showing a nuclear envelope and a nuclear pore complex (white arrow), and the tomogram segmentation (right) showing membranes (turquoise), nucleosome (yellow) and docked NPCs (pink, EMD-3103). Cyto=cytoplasm, nuc=nucleus. Scale bar: 100 μm. **(b)** Tomographic slice (left) showing a myelinated axon, and the tomogram segmentation (right) showing membranes (turquoise) and neurofilament (pink). Scale bar: 100 μm. **(c)** Tomographic slice (left) showing a synapse and its segmentation (right) showing membranes (turquoise) and F-actin filaments (right). a=axon terminal, s=synaptic cleft, d=dendrite. Scale bar: 100 μm. (**d**) Insets showing a synaptic vesicle fusing with the presynaptic membrane (top) and an actin filament in the postsynaptic dendrite (bottom, white arrows). a=axon terminal, d=dendrite. Scale bars: 50 nm. **(e)** Subvolume average of actin filaments (left) with EMD-15106 (pink) docket into it (right). Scale bar: 20 Å. **(f)** Tomographic slice (left) showing a neurite with neurofilament and its segmentation showing membranes (turquoise) and neurofilaments (pink). Scale bar: 100 nm. (**g**) Averaged 2D projections of extracted neurofilament subvolumes. Scale bar: 20 nm. **(h)** Reconstructed volume of neurofilament based on 2D-projected particles, implying a pentameric structure. Scale bar: 20 Å. **(i)** Subvolume average of neurofilaments (imposed symmetry based on vimentin intermediate filaments^32^).

**Extended Data Fig. 2:**
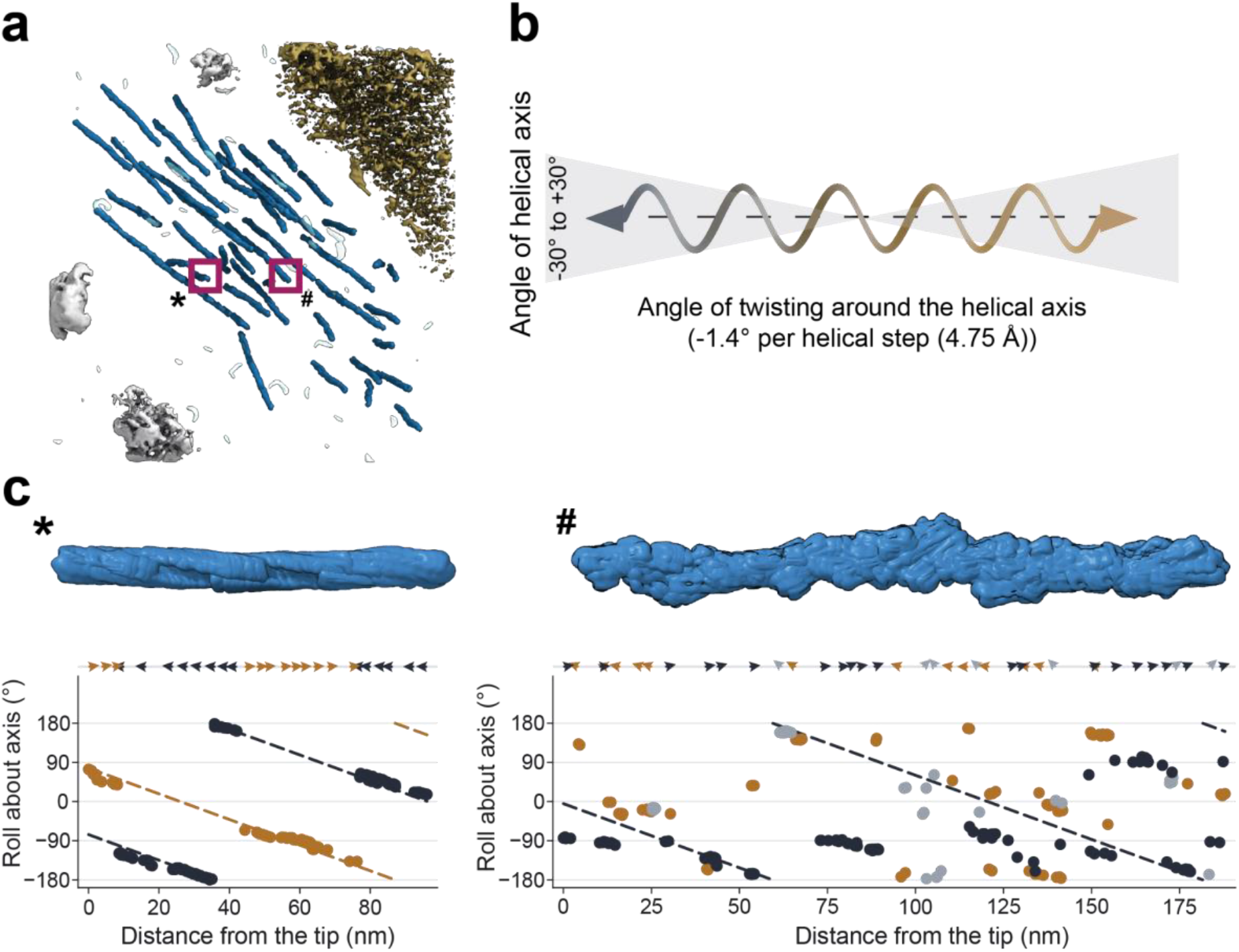
The Rohlex software to investigate helical reconstruction after 3D averaging. **(a)** A 3D rendering isosurface of MSA human brain tomogram. The magenta squares mark the filaments displayed in (c). **(b)** The scheme image to showcase the core equation of the toolbox. The refined poses of each segment of a filament are translated into 2 parameters: the angle of its helical axis in the 3D space and the helical twisting angle along the helical axis. The filament model axis was defined with 3D coordinates of all the segments (marked as double head dash line). The segment’s helical axis outside a cone range of 30° to model is marked as gray; the major polarities of the segments are marked as black and the antiparallel ones as brown. For α-syn filaments, a model parameter should have a -1.4° rotation along the helical axis for each helical step (4.75 Å). **(c)** Examples of a well aligned filament (*) and a badly aligned filament (#), selected from (a). The upper panel shows the isosurface rendering filament with the same helical axis pointing horizontally. The lower panel shows the polarity of each segment as colored arrowheads (same color scheme in (b)), and the scattered plot shows the relative rotation angle along the helical axis (0° at the center). The colored dash lines illustrate the model helix with slope of –1.4° per 4.75 Å.

**Extended Data Fig. 3:**
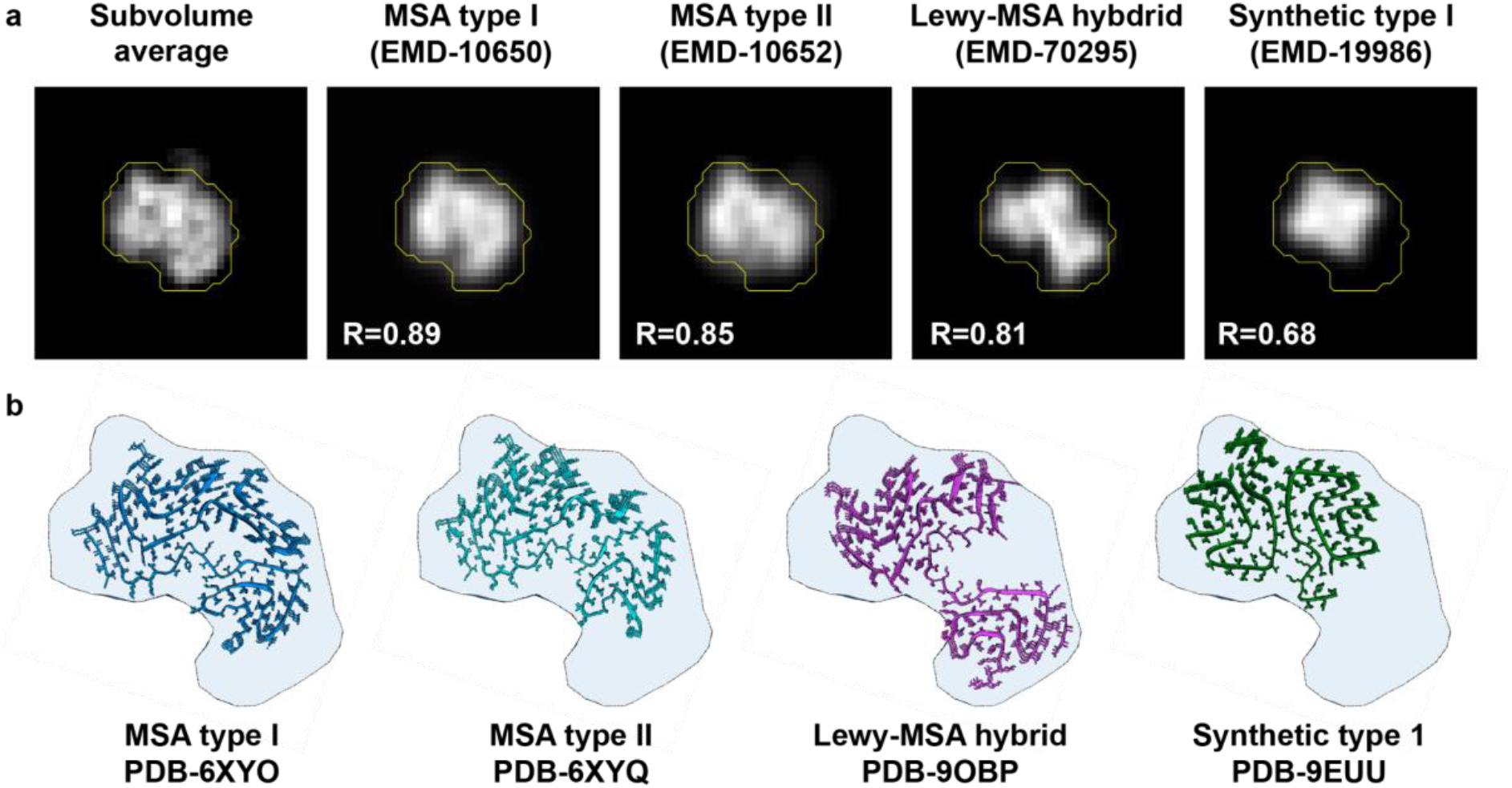
Comparison of the *in situ* α-synuclein subvolume average with disease-relevant double-protofilament structures. **(a)** Top views of the subvolume average (left) and maps of existing structures filtered to a similar resolution (30Å). The yellow outline indicates the mask region used to calculate the Pearson’s correlation coefficient (R), which was generated by thresholding the subvolume average and expanding it by 3 pixels. **(b)** Models of the corresponding atomic models fitted into the subvolume average.

**Extended Data Table 2:** specifics of the tilt series acquisition and subvolume averaging. Tau 1 refers to the density shown in Fig. 3c, tau 2 to the density shown in Fig. 3d, and α-syn to the density in Fig. 4c.

| | tau 1 | tau 2 | α-syn |
| --- | --- | --- | --- |
| Raw tilt series | EMPIAR-XXX | EMPIAR-XXX | EMPIAR-XXX |
| Reconstructed volume | EMD-XXX | EMD-XXX | EMD-XXX |
| **Data collection** |  |  |  |
| Nominal magnification | 53'000x | 53'000x | 64'000x |
| Pixel size (Å) | 2.42 | 2.42 | 1.98 |
| Voltage (kV) | 300 | 300 | 300 |
| Dose per tilt (e/Å<sup>2</sup>) | 3.78 | 3.78 | 3.8 |
| Tilt range | −50° to 70° | −50° to 70° | −50° to 67° |
| Tilt step (°) | 3° | 3° | 3° |
| Total dose (e/Å<sup>2</sup>) | 155 | 155 | 153 |
| Defocus range (μm) | −1 to −5 | −1 to −5 | −2.5 to −5 |
| **Subvolume averaging and helical reconstruction** |  |  |  |
| Box size | 64 | 64 | 64 |
| Binned pixel size | 9.68 | 9.68 | 7.92 |
| Initial subvolumes (no.) | 15'250 | 9'131 | 9'422 |
| Final subvolumes (no.) | 8'694 | 5'679 | 5'312 |
| Symmetry imposed | C1 | C1 | C1 |
| Helical twist (°) | −0.94 | −1.13 | −1.58 |
| Helical rise (Å) | 4.72 | 4.72 | 4.80 |
| Map resolution<br>(FSC threshold at 0.143) | 34 | 36 | 30 |

## Supplementary Information

**Supplementary video 1: Summary of the cryo-ET workflow applied to MSA postmortem brain.**

**<u>Supp-video-01.mp4</u>**

**Supplementary video 2: Tomographic volume and segmentation of a nuclear envelope and two nuclear pore complexes in MSA brain (shown in Extended Data Fig. 1a).** Membranes in turquoise, nucleosomes in yellow, a docked NPC (EMD-1506) in pink. Voxel size: 7.92 Å.

<u>Supp-Video-02.mp4</u>

**Supplementary video 3: Tomographic volume and segmentation of a myelinated axon in MSA brain (shown in Extended Data Fig. 1b).** Membranes in turquoise, neurofilament in pink. Voxel size: 7.92 Å.

<u>Supp-Video-03.mp4</u>

**Supplementary video 4: Tomographic volume and segmentation of a synapse in MSA brain (shown in Extended Data Fig. 1c).** Membranes in turquoise, actin filaments in pink. Voxel size: 7.92 Å.

<u>Supp-Video-04.mp4</u>

**Supplementary video 5: Tomographic volume and segmentation of a neurite in MSA brain (shown in Extended Data Fig. 1f).** Membranes in turquoise, neurofilament in pink. Voxel size: 7.92 Å.

<u>Supp-Video-05.mp4</u>

**Supplementary video 6: Tomographic volume and segmentation of tau filaments in a neurofibrillary tangle in AD brain (shown in Fig. 3c).** Membranes in turquoise, tau filaments in brown. Voxel size: 9.68 Å.

<u>Supp-Video-06.mp4</u>

**Supplementary video 7: Tomographic volume and segmentation of tau filaments in a neurofibrillary tangle in AD brain (shown in Fig. 3d).** Membranes in turquoise, tau filaments in brown. Voxel size: 9.68 Å.

<u>Supp-Video-07.mp4</u>

**Supplementary video 8: Tomographic volume and segmentation of a glial intranuclear inclusion in MSA brain (shown in Fig. 4b).** α-Syn fibrils in blue, nucleosomes in yellow, aggregates in grey. Voxel size: 7.92 Å.

<u>Supp-Video-08.mp4</u>

**Supplementary video 9: Tomographic volume and segmentation of a fibril crossing a glial nuclear envelope in MSA brain (shown in Fig. 5c).** Membranes in turquoise, mitochondrial membranes in red, α-syn fibrils in dark blue, nucleosomes in yellow. Voxel size: 7.92 Å.

<u>Supp-Video-09.mp4</u>

### Subtomogram averaging for *in situ* filaments in Cryo-ET

Subtomogram averaging requires multiple complex processing steps. So far, different software and platforms each serve specific advantages (RELION^25^, Scipion^45^, Dynamo^24^, Warp^27^ and emClarity^46^). Here, we present the state of the art STA for *in situ* amyloid filaments. For a step-by-step tutorial, see https://lbem-ch.github.io/amy-subtomo-averaging.

## 1. Preprocessing and data validation

To check tomography data from batch acquisition, a fast preprocessing approach must be undertaken to reconstruct tomograms at desired binned pixel size for image quality examination and tilt series-alignment. Here, we use the *tomotools*^39^ package for batch motion correction^47^ (*a*), tilt-series alignment with AreTomo^40^ and tomogram reconstruction with IMOD^30^ (*b*) at desired binned resolution (pixel size at 7.92 Å).

*A. tomotools preprocess --mcbin 1 --frames ./FrameFolder --gainref GainFile TiltSeries.mrc ./OutputFolder*

*B. tomotools reconstruct -b 4 -d 3000 --aretomo ./OutputFolder/CorrectedTiltSeries.mrc*

(Note: images acquired as Electron-Event Representation format (.eer) have to first be converted to movies of a chosen frame dose in .tif format with *relion_convert_to_tiff*)

The overall process took 5 to 10 mins per tomogram on a 120-CPU Linux computer with four NVIDIA A40 GPUs. The reconstructed tomogram at the binned resolution can then be validated with *3dmod* in IMOD. Tomograms with amyloid filaments of good quality were marked for further processing with Warp. For samples with gold fiducial markers, we recommend using IMOD semi-manual tilt-series alignment since AreTomo usually performs worse with high-contrast objects. *Tomotools* also enables importing/exporting alignment files from IMOD.

Due to the heterogenetic environment of native *in situ* samples, the amyloid filaments were preserved in various orientations within the acquired area (e.g., in **Fig. 3c,e**). Manual rendering of the dataset is usually unavoidable at this screening phase. We recommend properly marking down the rendered object in a separate datasheet.

The tomograms with amyloid filament and desired quality were then exported with *tomotools* for the Warp environment setup:

*C. tomotools aretomo2warp --v2 --frames-dir ./FrameFolder ./OutputFolder ./WarpFolder*

## 2. Prepare Warp/RELION/Dynamo environment

*Tomotools* exports the preprocessing files into 3 folders: *frames, imod,* and *mdoc*. The *imod* folder contains the alignment files in the IMOD format to be imported to *WarpTools*. To ensure the best alignment condition of the preprocessing steps, we recommend verifying the alignment quality with the aligned tilt-series _ali.mrc inside the *OutputFolder* from the previous step after export.

Generally, blurred images and images with objects blocking the view, *e.g.*, ice contamination or grid bars, should be excluded from processing in order to provide the most stable data quality. We recommend removing those images from the *FrameFolder*, editing the .mdoc file, and re-processing from the previous step.

*WarpTools* is then used to prepare config and header file settings for STA, including patch CTF estimation. Within our acquisition setup, the following commands can be executed with a shield script inside the *WarpFolder* folder:

*D. WarpTools create_settings --folder_data frames --output warp_frameseries.settings - -extension *.mrc --angpix 1.98 --exposure 3.83 --folder_processing processing*

*E. WarpTools fs_motion_and_ctf --settings warp_frameseries.settings --m_grid 1×1×1 -- c_grid 2×2×1 --c_range_min 50 --c_range_max 10 --c_defocus_max 8 -- out_averages --c_use_sum --out_average_halves*

*F. WarpTools ts_import --mdocs ./mdoc --frameseries ./processing --tilt_exposure 3.83 - -output ./processing/tomostar*

The above scripts create the basic settings for patch CTF estimation (D). Since motion-corrected images were exported in the previous steps, the motion grid (*--m_grid*) is set to *1×1×1* for no correction (E). The alignment information from *tomotools* can be directly imported to *WarpTools* (*F*). To improve the alignment results from AreTomo, MissAlignment is used, which is a deep learning based model within Warp^28^ (G). The following scripts set up the tilt-series config file (H), determined defocus handedness and estimate the overall tomogram CTF (*I* and *J*), and reconstruct the tomogram (*K*):

*G. miss-alignment --config-file /path/to/config.yaml --training-devices 0,1 -- reconstruction-devices 2,2,2,3,3,3 --dataloaders-per-trainer 5 --start-at-iteration 0 -- prepare-stacks 7.92*

*H. WarpTools create_settings --folder_data processing/tomostar --output warp_tiltseries.settings --extension *.tomostar --angpix 1.98 --exposure 3.83 -- folder_processing processing --tomo_dimensions 4096×4096×3000*

*I. WarpTools ts_defocus_hand --settings warp_tiltseries.settings --set_auto*

*J. WarpTools ts_ctf --settings warp_tiltseries.settings --defocus_max 8 --range_high 4*

*K. WarpTools ts_reconstruct --settings warp_tiltseries.settings --angpix 7.92 -- dont_invert --deconv*

The CTF-corrected tomogram is reconstructed inside the *WarpFolder/processing/reconstruction* folder and should be identical to the one reconstructed with *tomotools*. The *--deconv* flag (*K*) generates a denoised tomogram for easier segmentation. Since *WarpTools* generates linked config files, in case of any processing disruptions, the *.settings* files and the entire *processing* folder should be deleted to avoid conflicts from older versions. More information can be found in the detailed user manual for *WarpTools*^48^.

## 3. Segmentation

Tomogram segmentation is critical for STA. For amyloid filaments, it is crucial to identify not only the correct particles of interest but also their precise 3D coordinates and filament pose. The software package *IMOD* is therefore used to manually segment the amyloid filament from each chosen tomogram.

In *IMOD*, the program *3dmod* is used to pick filaments with individual contours of multiple points that follow the main stem of each filament. A filament should be picked continuously in order to measure the pre-align pose, which is crucial for initial model generation and alignment at the beginning of STA. The *IMOD* model is then saved as a *.mod* file. To generate equal and continuous segments, IMOD is used to generate segmentation with 7.92 Å interbox distance within a filament:

*L. addModPts ImodModel.mod 1*

This creates a new *_PtsAdded.mod* file from the enriched model. To generate a plain text file from the *IMOD .mod* file for later processing, IMOD function is called:

*M. model2point -c ImodModel_PtsAdded.mod ImodModel_PtsAdded.coords*

The last command creates a 4-column text file, in which each row records the filament-index and the x-y-z coordinates of each segment. This file *ImodModel_PtsAdded.coords* can then be used for creating the *.star* and *.coords* files for subtomogram reconstruction in *Warp*.

To create the *.star* file for *Warp* subtomogram reconstruction, *crYOLO* is used (*N*). Note that the input file for *crYOLO* has a different column order, in which the filament index should be at the last column. The first column of *ImodModel_PtsAdded.coords* therefore has to be moved to become the last. This creates a *particles_warp.star* file, containing the tilt and psi prior Euler angles and the filament tube ID:

*N. cryolo_boxmanager_tools.py coords2star -i ImodModel_PtsAdded_edited.coords -o . --apix 1.98 --mag 64000 --flipratio 0.5 --scale 4*

## 4. Subtomogram reconstruction

Subtomogram extraction is then done in *WarpTools* with GPU acceleration. The subtomograms are then further processed with *Dynamo* and *RELION* for 2D screening, 3D pre-alignment, and 3D refinement. These steps and their purposes are described below.

An important choice at each STA stage is thereby, if the workflow should manipulate the subtomograms as 3D volumes, or if only the coordinate parameters of sub-volumes should be refined and applied for fresh 3D reconstructions from the original tilt-series 2D image data.

In the first approach, each subvolume is treated as a small real-space 3D volume that has to be CTF-corrected with a 3D CTF mask that is prepared for each particle. In the second approach, as implemented in newer versions of RELION^31,49^, all image information is stored in the original tilt-series format to save storage space and computation time, and 3D reconstructions are computed from the raw image data.

In our pipeline, we follow both approaches, applying them at different stages, as described below. In *WarpTools*, both data types can be reconstructed with the flag --3D for 3D volume subtomogram (*O*) and --2D for tilt-series subtomogram handling (*P*) at the desired binned resolution. Here, bin8 is used as an example:

*O. WarpTools ts_export_particles --settings warp_tiltseries.settings --input_star particles_warp.star --output_star OutputFolder_3D particles.star --coords_angpix 1.98 \ --output_angpix 15.84 --box 64 --diameter 1013 --relative_output_paths **--3d** -- output_processing SubtomogramFolder_3D*

Or alternatively:

*P. WarpTools ts_export_particles --settings warp_tiltseries.settings --input_star particles_warp.star --output_star OutputFolder_2D particles.star --coords_angpix 1.98 \ --output_angpix 15.84 --box 64 --diameter 1013 --relative_output_paths **--2d** -- output_processing SubtomogramFolder_2D*

Note that *particles_warp.star* might contain the wrong data name for the header

*_rlnMicrographName*. The data name for the *_rlnMicrographName* should be corrected to be the name of *.tomostar* files generated from Warp during processing. This is a syntax error using *cryolo_boxmanager_tools.py* for *WarpTools*.

## 5. Subtomogram averaging

### 2D screening in RELION (optional step)

Subtomogram averaging suffers from missing wedge artifacts. To get the first idea of how the data look like, one of the compromised approaches is to project or average the CTF-corrected 3D volumes along the electron gun axis into 2D images, and follow the standard single particle analysis (SPA) pipeline, including 2D classification, initial model generation, and 3D refinement ^50,51^. The approach serves as an optional quality control for the coordinate and systematic alignment. This step can be skipped when the user is confident about the data quality.

After subtomogram reconstruction with *(O)*, the 3D volumes were then corrected by multiplying its CTF mask (*CTF_volume*), and projected along the image Z-axis, masked with desired thickness:

*Q. CTF_corrected_volume = IFFT(FFT(subtomogram_volume*CTF_volume))*

*R. projection(x,y) = mean(z)(CTF_corrected_volume(x,y,z))*

The projection images can be used for 2D classification in RELION. Due to the dimension reduction, one can use the unbinned pixel size and a bigger box size in (*O*) to have a more comprehensive view of the filament if necessary. Notably, depending on the Warp version, the output star file might need additional modification, such as adding

*_rlnHelicalTrackLengthAngst*, *_rlnAnglePsiFlipRatio*, and *_rlnHelicalTubeID*, for the helical reconstruction. The selected 2D classes and particles could then be directly used for 3D refinement in RELION or as screening index for 3D subtomogram averaging in Dynamo.

#### Dynamo

Here we used Dynamo to initiate bias-free 3D subtomogram averaging, prior to structural refinement in RELION. We first imported the RELION/Warp star file to the Dynamo particles.tbl file with *warp2dynamo* ^52^ (*S*). The particles.tbl file contains the corresponding particle prior-pose information for the initial model generation.

*S. warp2dynamo -i path_to_star_file -o path_to_dyanmo_project/particles -bs 64*

In dynamo, particles are read as dark value .em files, which are inverted to Warp’s output. As a result, after CTF correction (Q), we used the EMAN2 package for conversion. ^53^ (*T*).

*T. e2proc3d.py --mult=-1 CTF_corrected_volume.mrc output.em*

The output.em files were named numerically based on the first column of the particles.tlb file (e.g. 1.em, 2.em) , and should be stored in the same folder. In our setting, the particle’s axis was rotated in the default of Dynamo. As a result, the tilt axis was set to the x-axis and the range was set to minus. For example, in the general setup, a tomogram acquired by a scheme of -50° to 70° of y-axis, we marked the tomogram as -70° to 50° of x-axis in dynamo. We then edited the particles.tbl with the 13th column to 2, and the 15th and 16th columns as -70 and 50. This is important for generating the missing wedge Fourier mask during particle alignment in Dynamo.

Dynamo can be executed with MATLAB Compiler Runtime (MCR) in Ubuntu terminal with installing MATLAB. The user can run the Dynamo command lines within the Dynamo console. We started the subtomogram averaging from 8x binning to improve efficiency. First, we generated the featureless initial reference by randomly rotating the particles along its filament axis. This approach can also reduce the effect of missing wedge trap:

*In Dynamo console:*

*T = dread(’particles_edit.tbl’); T2 = dtrandomize_azimuth(T);*

*dwrite(T2,’particles_edit_mod.tbl’); oa=daverage(’particle_folder’,’t’,’particles_edit_mod.tbl’,’mw’,50); dview(oa.average);*

*dwrite(oa.average,’raw_template.em’);*

The Dynamo project named bin8_align can then be launched with the generated reference,

*raw_template.em*, and table, *particles_edit_mod.tbl*:

*dcp.new(’bin8_align’, ’d’, ’particle_folder’,’template’,’raw_template.em’,’masks’, ’default’,’t’,’particles_edit_mod.tbl’);*

The Dynamo project GUI will show. In the GUI, the mask tool can generate the desired cylinder mask for the alignment job. For this step, we used a mask covering around 80% of the particle size to allow the particle to align better along the filament axis. This mask is used for alignment, classification, and FSC calculation. The Fourier template mask is set to one as no mask.

In the numerical parameters, Dynamo allows users to specify alignment search schemes. Since the prior pose has been determined during segmentation, we limited alignment search as follow:

| Round 1 | Round 2 |
| --- | --- |
| Iterations: 2 | Iterations: 2 |
| Cone aperture: 20 | Cone aperture: 20 |
| Cone sampling: 3 | Cone sampling: 3 |
| Cone flip (Advanced): 1 | Cone flip (Advanced): 1 |
| Azimuth rotation range: 360 | Azimuth rotation range: 45 |
| Azimuth rotation sampling: 15 | Azimuth rotation sampling: 5 |
| Refine: 2 | Refine: 2 |
| Low: 25 | Low: 25 |
| Shift limits: 4 4 2 | Shift limits: 4 4 2 |
| Shift limiting way: 4 | Shift limiting way: 4 |

In the computing environment, we used the 4 GPUs in the GPU standalone mode with 50 MATLAB pool engines. Use the check and unfold buttons to generate the *bin8_align.exe*, which can be executed in the terminal.

In our case, at the 4th iteration of this project, the average 3D volume (*average_ref_001_ite_0004.em*) had already allowed us to see and determine the helical parameters. We used the Rohlex software we developed to determine the helical rise and twist (**Extended Data Fig 1**), and relion_helcial_toolbox to impose the symmetry. The imposed volume (*average_ref_001_ite_0004_sym.em*) was then used for the next refine run in Dynamo.

Next, we ran the adaptive bandpass project in Dynamo:

*dcp.new(’abp_align’, ’d’, ’particle_folder’,’template’, ’average_ref_001_ite_0004_sym.em’,’masks’,’default’,’t’,’refined_table_ref_00 1_ite_0004.tbl’);*

This time, we used a smaller cylinder mask that covered only around 50% of the box size. Set the numerical parameters with only 1 round: Iter 4, cone flip 2, azymuth rotation range 180, azymuth rotation sampling 15, azymuth flip 2, refine 3, low 25, symmetry h[-twist,rise]. Notably, as we mentioned before, the tilt axis is recognized as flipped in Dynamo, the helical twist is therefore mirrored as *-twist*.

In the GUI, press Multiference, Adaptive filtering (“golden standard”), and Derive a project. This will generate the golden standard refinement job (abp_align_eo) comparing the average of even/odd particles. Point the project name to abp_align_eo to open the project. In the Multiference, Adaptive filtering (“golden standard”), and Edit parameters for Adaptive Filtering run, edit the adaptive bandpass threshold to 0.143 and the adaptive bandpass pushback to 0. Unfold and run the job.

The output of the abp_align_eo can be investigated with the Rohlex software. The filaments that did not follow the determined helical parameters were removed for the next job.

Next, we used the output at 8x binning to generate the star file in Warp to reconstruct the subtomogram at 4x binning. The conversion from Dynamo to Warp can be done with *dynamo2warp* ^52^ (*U*):

*U. ’dynamo2warp’,’-i’,dynamo_table.tbl,’-tm’,’particles.reextract.doc’,’-o’, ’particles.star’*

At 4x binning, we repeated the same workflow of subtomogram reconstruction, CTF correction, and creating the Dynamo project (*O*, *Q, S, and T*). The reference for the alignment can also be created with the same Dynamo command at 8x binning. We created the adaptive bandpass project at 4x binning and performed the alignment with helical symmetry, with the same numerical parameters at 8x binning.

In our cases, the filament polarity can be observed from the averaged structures. To further improve the alignment outcome, we used the Rohlex software to normalize polarity by flipping misaligned segments, and created a new fixed Dynamo table for another adaptive bandpass project with cone flip = 0. This approach forced the alignment to focus on the correct filament polarity.

Considering the low S/N in *in situ* Cryo-ET data, the outcome of our Dynamo projects established the better initial particle poses and cleaner dataset that are crucial for the Bayesian approach refinement in RELION.

#### RELION

We then convert the Dynamo output at 4x binning to Warp and RELION by using the commands (*U*). Since RELION version 5 supports 2D tilt-series subtomogram averaging, we use the (*P*) to extract the 2D subtomogram and to speed up the processing. The reference volumes were generated from the Dynamo command and inverted to the .mrc format. The masks for RELION jobs were generated by the *relion_helical_toolbox*.

In RELION, the initial particle poses of helical reconstruction are defined with the star file headers *_rlnTomoSubtomogramRot/Tilt/Psi*. These angles can be properly handled by subtracting a 90° pre-rotation from the outcome from Dynamo refinement in *Tilt angle*. In our methods, we also added the *_rlnCenteredCoordinateX/Y/ZAngst*, *_rlnAngleRot/Tilt/PsiPrior*, *_rlnAnglePsiFlipRatio*, *_rlnHelicalTubeID*, and *_rlnHelicalTrackLengthAngst* to allow the full functions for helical reconstruction in RELION.

We then performed a 1 class 3D classification (K = 1) in RELION with helical reconstruction with T = 4 and limited Limit resolution E-step = 20 to further improve the alignment with limited particles. The badly aligned particles were removed with the invest_helical toolbox to reduce overfitting. Finally, the output was used for a 3D refine job for the gold-standard refinement to confirm the resolution.

Depending on the data conditions and users’ needs, the refined output of RELION, *run_data.star*, could be used to reconstruct the 2D subtomograms (*P*) at the finer pixel sizes and bigger box size. The 2D subtomogram approach in RELION allows fast 3D refinement and classification even with large datasets, to ensure the efficient trial-and-error iteration cycle.

## Notes

https://lbem-ch.github.io/amy-subtomo-averaging

https://github.com/LBEM-CH/rohlex

### Competing Interest Statement

The authors have declared no competing interest.

